# Partitioning and Enhanced Self-Assembly of Actin in Polypeptide Coacervates

**DOI:** 10.1101/152025

**Authors:** Patrick M. McCall, Samanvaya Srivastava, Sarah L. Perry, David R. Kovar, Margaret L. Gardel, Matthew V. Tirrell

## Abstract

Biomolecules exist and function in cellular micro-environments that control their spatial organization, local concentration and biochemical reactivity. Due to the complexity of native cytoplasm, the development of artificial bioreactors and cellular mimics to compartmentalize, concentrate and control the local physicochemical properties is of great interest. Here, we employ self-assembling polypeptide coacervates to explore the partitioning of the ubiquitous cytoskeletal protein actin into liquid polymer-rich droplets. We find that actin spontaneously partitions into coacervate droplets and is enriched by up to ≈30-fold. Actin polymerizes into micrometer-long filaments and, in contrast to the globular protein BSA, these filaments localize predominately to the droplet periphery. We observe up to a 50-fold enhancement in the actin filament assembly rate inside coacervate droplets, consistent with the enrichment of actin within the coacervate phase. Together these results suggest that coacervates can serve as a versatile platform in which to localize and enrich biomolecules to study their reactivity in physiological environments.

**SIGNIFICANCE STATEMENT:** Living cells harbor many protein-rich membrane-less organelles, the biological functions of which are defined by compartment composition and properties. Significant differences between the physico-chemical properties of these crowded compartments and the dilute solutions in which biochemical reactions are traditionally studied pose a major challenge for understanding regulation of organelle composition and component activity. Here, we report the spontaneous partitioning and accelerated polymerization of the cytoskeletal protein actin inside model polypeptide coacervates as a proof-of-concept demonstration of coacervates as bioreactors for studying biomolecular reactions in cell-like environments. Our work introduces exciting avenues for the use of synthetic polymers to control the physical and biological properties of bioreactors in vitro, enabling studies of biochemical reactions in cell-like micro-environments.

## INTRODUCTION

The biological functions of intracellular organelles are defined by the composition and properties of the compartments, which often differ significantly from that of bulk cytoplasm. Well known examples include the acidic pH of lysosomes and the mitochondrial redox potential (1). While the compartmentalization of these organelles require a lipid bilayer as a physical barrier, recent work has shown that organelles can also form as phase-separated droplets that do not require such a membrane (2, 3). The physicochemical properties of membrane-less organelles likely regulate partitioning and reactivity of biomolecules, thereby serving an important role in their physiological function. The compositional complexity of individual cellular bodies, granules, and organelles pose a major challenge in discerning general mechanisms for partitioning and reaction regulation. One useful strategy has been to reduce compositional complexity by *in vitro* reconstitution of cellular bodies (4, 5). However, the sequence and structural complexity of natural biopolymers make systematic variation of micro-environment properties difficult.

A complementary approach is to selectively tune the physical and chemical properties of phase-separated micro-environments through the rational design of synthetic polymers that spontaneously phase separate via known mechanisms, and then use these materials as a platform to study biomolecule partitioning and reactivity. For instance, charged homopolymers (polyelectrolytes) form polymer-dense liquid phases via complex coacervation (6, 7) and localize charged proteins (8–10). Precise chemical control of polypeptide-based polyelectrolytes allows for fine-tuning of several physio-chemical properties of the coacervate phase (7, 11), including functional groups, water content, viscosity, and surface tension, thereby enabling systematic investigations of protein interactions and activities in controlled micro-environments (12, 13). Knowledge of the general mechanisms by which micro-environment properties tune protein partitioning and activity could provide needed insight into the function of membrane-less organelles as well as design principles for synthetic biology and engineering applications.

Here, we report the spontaneous partitioning and polymerization of the cytoskeletal protein actin inside model polypeptide coacervates (14, 15) as a proof-of-concept demonstration of coacervates as bioreactors for studying biomolecular reactions in cell-like physical environments. Our results establish polyelectrolyte complex coacervates as a viable platform to study mechanisms of partitioning and biochemical regulation by controlled perturbation of condensed-phase micro-environment.

## RESULTS

We use a model coacervate system (15) composed of the polycation poly-L-lysine (pLK) and the polyanion poly-(L,D)-glutamic acid (pRE), typically with ~100 amino acids per polypeptide (see Supplementary Materials and Methods, Table S1). Phase separation at room temperature is rapid; initially clear aqueous solutions become visibly turbid in seconds upon mixing of pLK- and pRE-containing solutions at total polypeptide concentrations of 10 μM or more (Movie S1), and is driven primarily by the release of condensed counterions (16). The presence of a polydisperse size distribution of polypeptide-rich coacervate droplets in solution, ranging in size from ~0.4 < R < 4 μm, is confirmed directly by differential interference contrast (DIC) microscopy (Fig. 1A-C, Fig. S1). The round, droplet-like appearance of the condensed pLK/pRE coacervate phase is suggestive of a fluid phase (15). Under similar conditions, the surface tension has been measured to be γ ~ 1 mN/m (17). Consistent with liquid-like properties on the timescale of seconds and longer, merging pLK/pRE droplets rapidly coalescence into a single, larger droplet (Fig. S1). From coalescence observations, we estimate the inverse capillary velocity *v*^−1^ = *η/γ* = 1.6 ms/μm (Fig. S1, (5, 18)). This yields a viscosity of η = 1.6 Pa·s, ~1000-fold higher than water. Thus, this simple model system is sufficient to create viscous phase-separated droplets with picoliter volumes.

**Figure 1.**
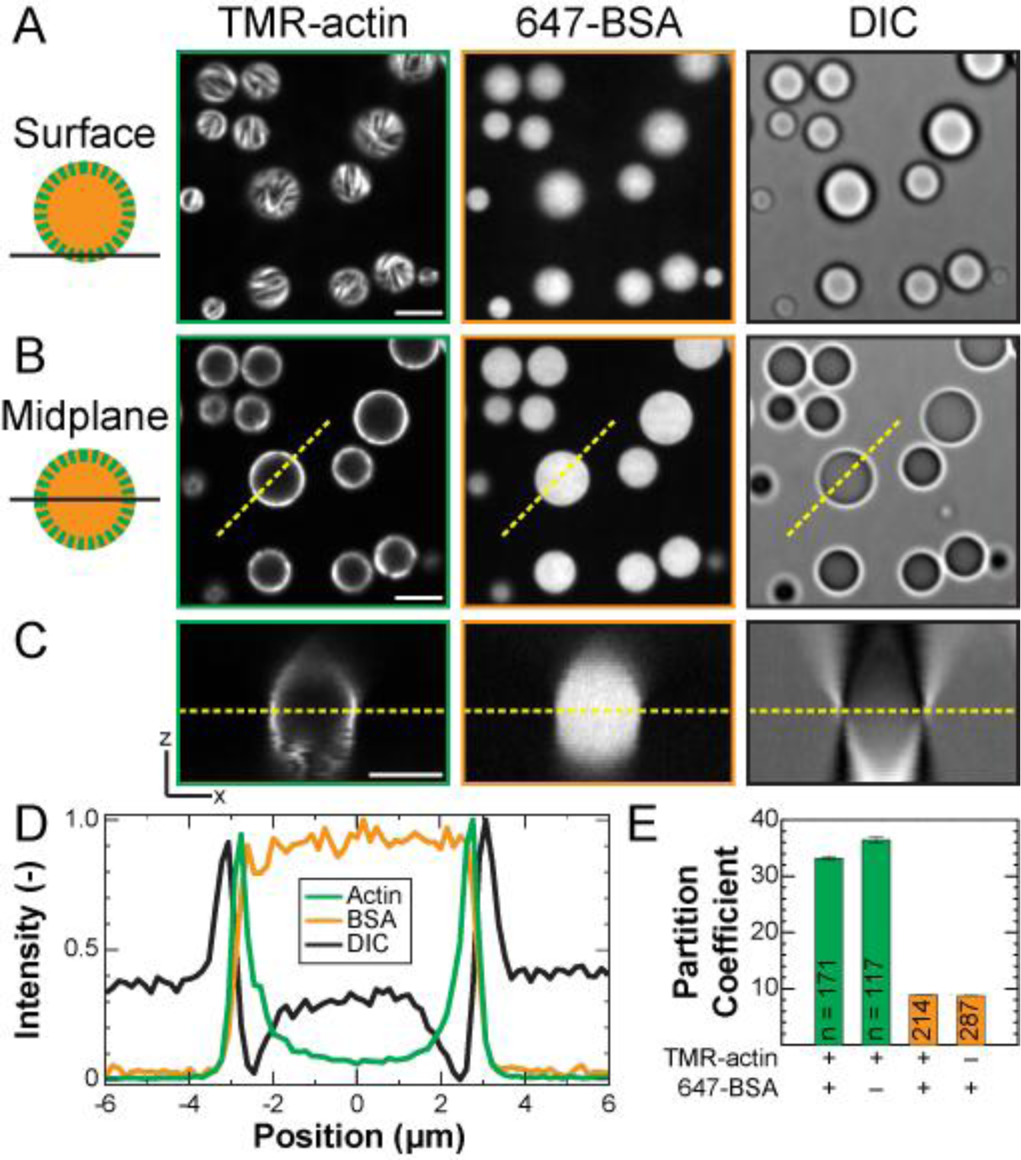
F-actin localize to the periphery of polypeptide coacervates. (A-B) Confocal fluorescence (left and middle) and DIC (right) micrographs of polypeptide coacervates containing both TMR-actin (green) and 647-BSA (orange) on non-adherent substrates. (A) is focused at the interface of the coacervates and the substrate (surface), and (B) is approximately the droplet midplane, indicated by the dashed yellow line in (C). (C) x-z cross-section taken along the dashed yellow line in (B) applied to all planes of a confocal z-stack. Scale bar is 5 μm in (A-C). (D) Normalized intensity linescans along the dashed yellow lines indicated in (B-C). (E) Average partition coefficient for coacervates containing 0.5 μM actin alone, 0.5 μM BSA alone, or 0.25 μM actin and 0.25 μM BSA (A-D). Error bars denote standard error of the mean. The number of droplets included in each condition is listed on the bar. Conditions are 0.5 μM total protein (0.25 μM Mg-ATP-actin (47% TMR-labeled) and 0.25 μM BSA (91% Alexa-647-labeled)) incubated with 5 mM pLK prior to addition of 5 mM pRE in 50 mM KCl,1 mM MgCl_2_, 1 mM EGTA, 10 mM imidazole (pH 7.0), and 72 μM ATP (all concentrations final).

### Charged proteins spontaneously partition into coacervate droplets

Using a previously published protocol, proteins are mixed with the cationic pLK prior to initiation of phase separation by the addition of anionic pRE (8). It was previously found that the negatively-charged protein BSA localizes preferentially to pLK/pRE coacervates, and is uniformly distributed within them (Fig. 1A-D, Fig. S2). This preference for the coacervate phase is described quantitatively by a partition coefficient, defined as the ratio of fluorescence intensity inside to outside the coacervates (4). We find an average partition coefficient of PC_avg_ ≅ 8, whether BSA is added to solution prior to or following phase separation (Fig. 1E, Fig. S3), indicative of spontaneous partitioning.

Here, we study the partitioning of actin, a cytoskeletal protein that self-assembles to form linear filaments (F-actin). Actin monomers and the chemically inert BSA are globular proteins of similar size (42 and 66 kDa, respectively) and carry comparable negative charge (isoelectric points of 5.23 and 5.60) (19). We find that actin partitions to pLK/pRE coacervates and immediately observe linear structures localized preferentially to the coacervate periphery (Fig. 1A-D). Integrating the total actin intensity within the droplet, we find an average partition coefficient that is 4-fold higher than that for BSA (Fig. 1E). Interestingly, the partition coefficients for BSA and actin are the same whether one or both proteins are present in solution (Fig. 1E, Fig. S2). This suggests that, under the conditions explored here, BSA and actin do not compete directly for space in the coacervate. Both partitioning and peripheral localization of actin are robust to the order of addition (Fig. S4).

### Self-assembled F-actin of canonical structure localizes to the coacervate periphery

To test whether the linear actin structures are bona fide F-actin, we stained with fluorescently-labeled phalloidin (647-phalloidin). Phalloidin is a small, uncharged toxin recognized for its ability to specifically bind to F-actin (20). 647-phalloidin was introduced into the solution after the coacervate formation and actin assembly, and found to localize along the linear actin structures. Confocal fluorescence micrographs at both the coverslip surface (Fig. 2A,C) and droplet midplane (Fig. 2B,D) reveal strong co-localization of phalloidin fluorescence to the linear actin structures with a Pearson’s correlation coefficient of 0.86 (Fig 2E, Supplementary Materials and Methods). This provides strong evidence that these linear actin-rich structures are composed of F-actin of canonical structure. Given the brightness of the F-actin structures, and previous work demonstrating that concentrations of polycations (and pLK in particular) lower than those in the coacervate phase are sufficient to bundle F-actin (21), we presume that the structures visible in Figs. 1 and 2 are F-actin bundles, rather than individual filaments.

**Figure 2.**
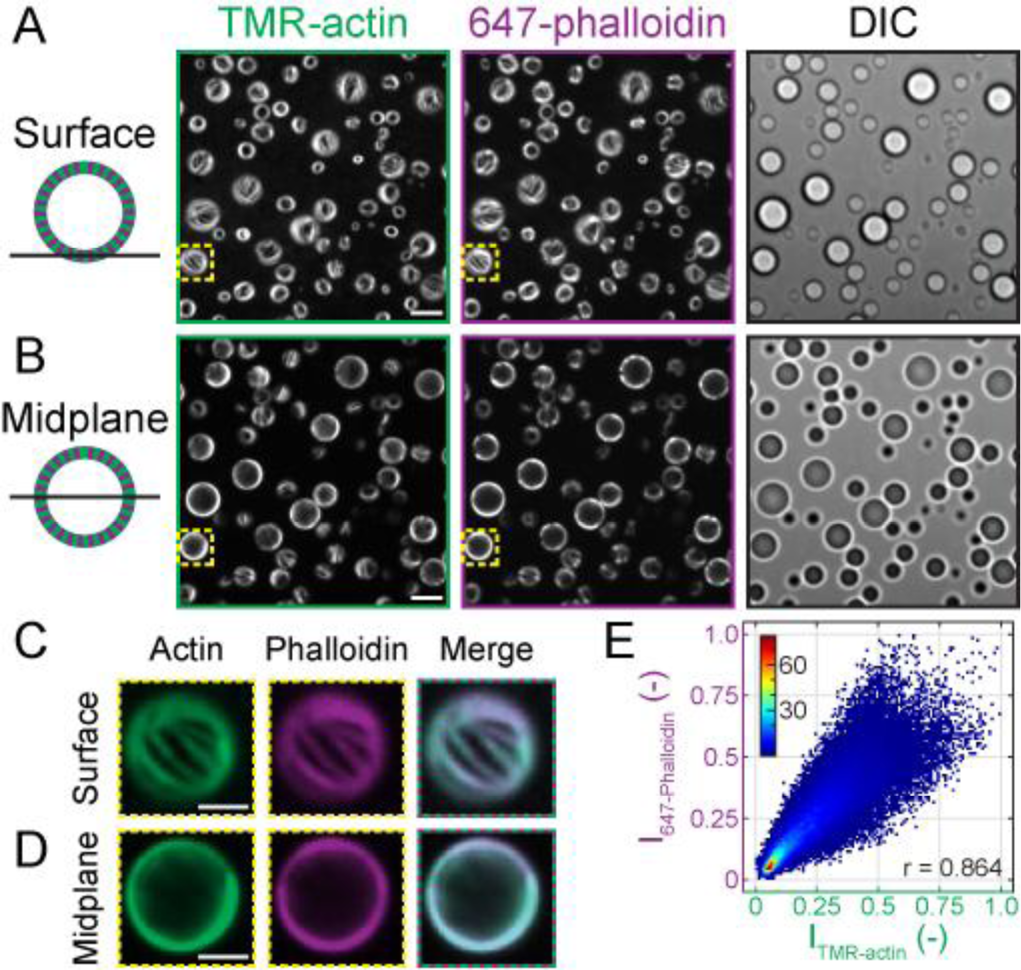
Linear actin fibers maintain canonical F-actin structure. (A-B) Confocal fluorescence (left and middle) and DIC (right) micrographs of polypeptide coacervates containing TMR-actin (green) after the addition of Alexa-647-Phalloidin (purple) on non-adherent substrates. (A) is focused at the interface of the coacervates and the substrate (surface), and (B) is approximately the droplet midplane. Scale bar is 5 μm. Conditions are 0.5 μM Mg-ATP-actin (47% TMR-labeled) incubated with 5 mM pLK prior to addition of 5 mM pRE in 50 mM KCl, 1 mM MgCl_2_, 1 mM EGTA, 10 mM imidazole (pH 7.0), and 72 μM ATP (all concentrations final). 0.25 μM Alexa-647-Phalloidin was flown into the chamber in the same buffer after droplets had sedimented. (C-D) False-colored fluorescence images of the regions outlined in yellow boxes in (A-B) from the surface (C) and midplane (D). Right column shows a merge. Scale bar is 2 μm. (D) Correlation between TMR-actin and 647-phalloidin fluorescence intensity values for all pixels in (A). Colors represent count density. Pearson’s correlation coefficient is r = 0.864.

### Actin assembly is enhanced in coacervates

Having demonstrated that the actin polymerization proceeds in pLK/pRE coacervates, we next ask to what extent the coacervate micro-environment impacts the reaction rate. Actin is a convenient model protein for this purpose owing to the existence of established spectroscopic tools for quantitatively monitoring assembly kinetics (22). In particular, the fluorescence intensity of the pyrene fluorophore increases ~20-fold when pyrene-labeled monomers are incorporated into filaments and is a well-established method to track actin assembly (22), as depicted in the schematic in Fig. 3A. In solution, the polymerization time course of 1.5 μM actin shows a characteristic lag phase, indicative of the kinetically slow filament nucleation step (23), followed by a phase of rapid growth and then saturation once a steady-state is reached (Fig. 3A) (22). At this actin concentration, the initial lag phase is typically ~10 min and steady-state is reached in ~120 min (Fig. 3B, black). The presence of pLK/pRE coacervates eliminates the lag phase and steady-state is achieved within 10 minutes (Fig. 3B, red). Thus, actin filament assembly is stimulated significantly by pLK/pRE coacervates.

**Figure. 3.**
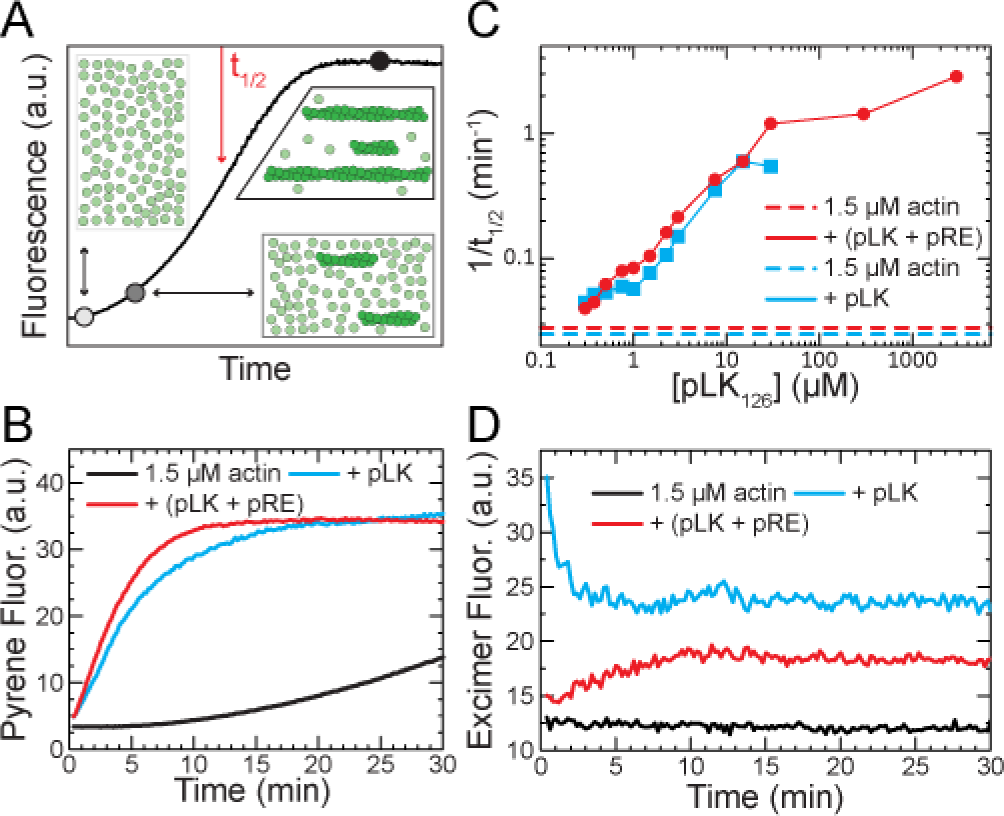
Coacervates and poly-L-lysine enhance actin assembly via different mechanisms. (A) Cartoon depicting the timecourse for spontaneous actin assembly monitored by changes in pyrene-actin fluorescence. (B) Spontaneous assembly of 1.5 μM Mg-ATP-actin (12% Pyrene-labeled) alone (black), with 3 μM pLK (cyan), or with 3 μM pLK and 3 μM pRE (red). (C) The assembly rate (1/t_1/2_) for 1.5 μM actin for samples with pLK alone (cyan) or equal concentrations of pLK and pRE (red) as a function of the concentration of pLK. Dashed lines denote the assembly rate of 1.5 μM actin alone measured in parallel with pLK-containing (cyan dashed) or pLK- and pRE-containing (red dashed) samples. (D) Timecourse of pyrene excimer fluorescence during spontaneous assembly of 1.5 μM Mg-ATP-actin (12% Pyrene-labeled) alone (black), with 3 μM pLK (cyan), or with 3 μM pLK and 3 μM pRE (red). In all experiments with polypeptides, Mg-ATP-actin is incubated with variable pLK in low salt prior to addition of pRE (red) or a buffer blank (cyan) in 50 mM KCl, 1 mM MgCl_2_, 1 mM EGTA, 10 mM imidazole (pH 7.0), and 72-150 μM ATP (all concentrations final).

To assess reaction kinetics quantitatively, we estimate the assembly rate 1/*t*_1/2_, defined as the inverse of the time at which the pyrene fluorescence intensity reaches half of its relative change during the course of actin polymerization (Fig. 3A, Supplementary Materials and Methods). The actin assembly rate 1/*t*_1/2_ increases from 0.03 to >1 min^-1^ as the total pLK concentration increases from 0.3 to 30 μM, while maintaining a pLK:pRE ratio of 1 (Fig. 3C). Above 30 μM, the assembly rate saturates. Thus, the actin assembly rate is enhanced by nearly two orders of magnitude in the presence of coacervates (Fig. 1, 2, 4).

### Polylysine and coacervates stimulate actin assembly via distinct mechanisms

One possible explanation for the enhanced assembly rate is polycation-mediated F-actin nucleation. Polylysine has been shown to promote formation of antiparallel actin dimers (24) that nucleate F-actin (25, 26). Spontaneous assembly of pyrene-labeled actin in the presence of pLK shows a concentration-dependent increase in the rate of actin assembly (Fig. 3B,C, blue data). It is tempting to compare the filament formation rate in solution directly to the rates observed within coacervates. However, the local pLK concentration within the coacervate phase is actually much higher, on the order of 1-3 M, such that the pLK concentrations reported in Fig. 3C should not be directly compared. Furthermore, pLK/pRE interactions within the coacervate could limit pLK-mediated antiparallel dimer formation.

To test whether pLK-stabilized antiparallel actin dimers contribute to the assembly of F-actin in pLK/pRE coacervates, we monitored pyrene excimer fluorescence (24). In the absence of pLK, 1.5 μM actin displays no change in pyrene excimer fluorescence during the nucleation-dominated early phase of assembly (Fig. 3D, black). In the presence of pLK, excimer fluorescence is highest during the initial nucleation phase, and decays rapidly as assembly proceeds (Fig. 3D, blue). This excimer fluorescence time course is the hallmark of actin assembly mediated by pLK-stabilized antiparallel actin dimers (24). Importantly, in the presence of pLK/pRE coacervates, excimer fluorescence does not have these features characteristic of anti-parallel dimer-mediated nucleation events (Fig. 3D, red). These data strongly suggest that pLK-mediated nucleation is not the dominant mechanism by which actin assembly is enhanced in pLK/pRE coacervates.

### Partitioning increases the local protein concentration in coacervates

A direct consequence of partitioning is that the local actin concentration in the coacervate phase is higher than that in the polymer-dilute phase. Thus, an alternate mechanism underlying enhanced assembly rates is an increased local actin concentration, c_local_, within coacervates.

We tested this possibility by varying the global actin concentration, c_global_, from 0.01 μM to 1.5 μM. The threshold monomer concentration, or critical concentration c*, required for polymerization of Mg-ATP-actin is ≈0.1 μM (27, 28). If actin is concentrated in coacervate droplets ≈30-fold via partitioning, we would expect actin assembly within coacervates at global actin concentrations of ≈0.003 μM. Importantly, we observe coacervate-associated F-actin at global actin concentrations as low as 0.05 μM (Fig. 4A-C). Interestingly, we observe peripherally-biased partitioning of actin to pLK/pRE coacervates at all actin concentrations examined, even at the lowest concentration (0.01 μM) for which no filaments are clearly discernible. We note that the density of peripherally localized F-actin changes as a function of the global actin concentration. Whereas isolated filaments or bundles are visible at 0.05 μM, an F-actin shell too dense to resolve individual structures forms at 1.5 μM (Fig. 4A-C).

**Figure 4.**
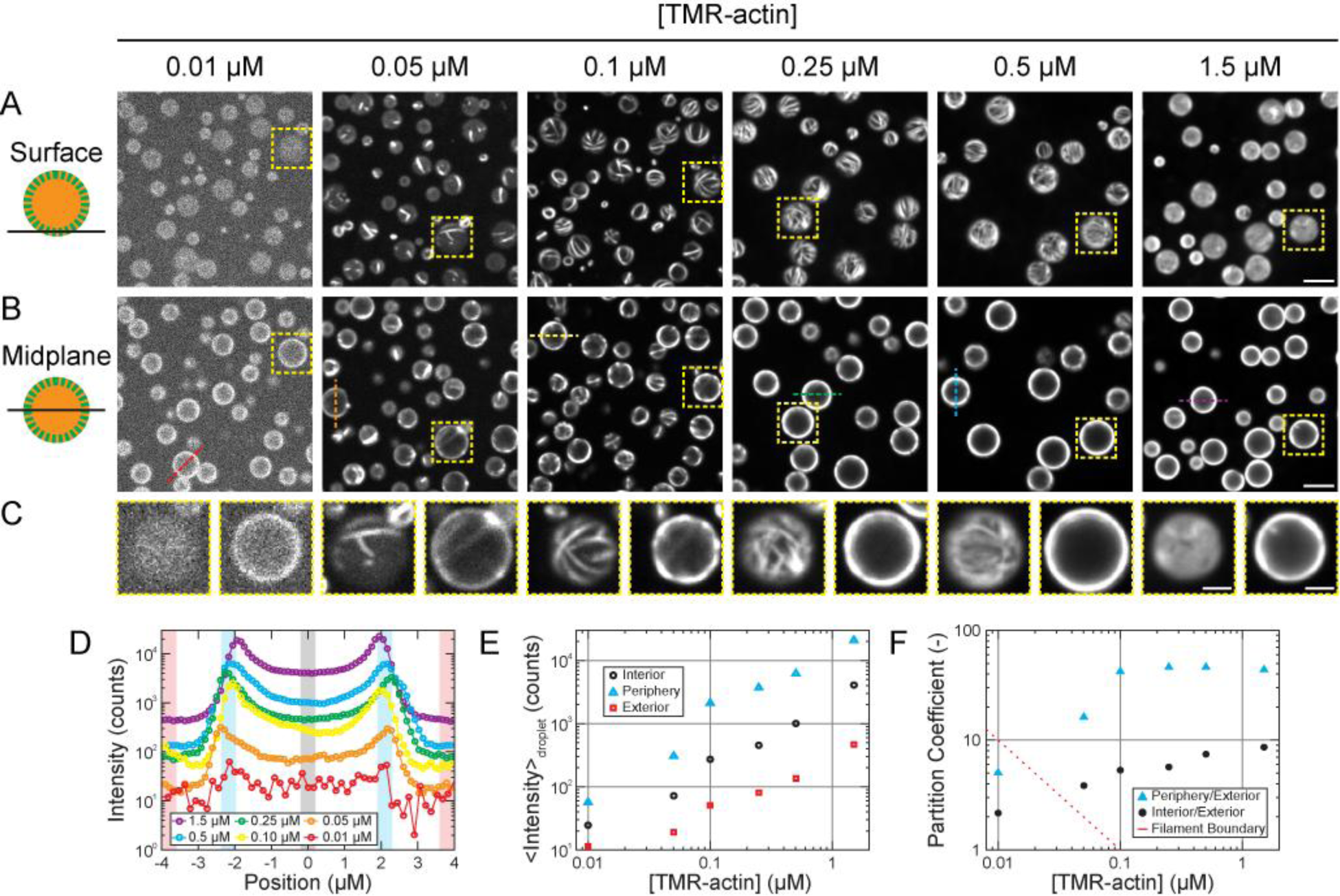
Partitioning increases the local protein concentration in coacervates. (A-B) Confocal fluorescence micrographs of polypeptide coacervates containing variable concentrations of TMR-actin on non-adherent substrates, focused at the interface of the coacervates and the substrate (surface, A), or approximately the droplet midplane (B). Contrast is adjusted individually for each concentration of TMR-actin, but is consistent between confocal slices for each condition. (C) Magnified images of the yellow boxed regions outlined in (A-B). Scale bars are 5 μm (A-B) and 2 μm (C). (D) Fluorescence intensity along the colored dashed lines in (B). (E) Mean fluorescence intensity of interior (circle, black), periphery (triangle, cyan), and exterior (square, red) of coacervates droplets, obtained from the black-, cyan-, and red-shaded regions in (D). (F) Partition coefficients (ratio of peripheral to exterior (triangles, cyan) and interior to exterior (circles, black) fluorescence) for the data in (D-E). Red line indicates where product of the partitioning coefficient and the total actin concentration equals the critical concentration for actin assembly of ~0.1 μM. Filaments are expected to the right of the red line, but not to the left. Conditions are a range of Mg-ATP-actin (47% TMR-labeled) concentrations incubated with 5 mM pLK (either alone (1.5- and 0.5-μM actin) or with 0.25 μM or BSA (91% Alexa-647-labeled) (0.25-, 0.1-, 0.05-, 0.01-μM actin)) prior to addition of 5 mM pRE in 50 mM KCl, 1 mM MgCl_2,_ 1 mM EGTA, 10 mM imidazole (pH 7.0), and 72 μM ATP (all concentrations final).

To systematically characterize the localization of actin fluorescence, we examined fluorescence intensity line scans through the midplane, depicted in Fig. 4B and shown in Fig. 4D. These curves may be divided into three regions: the droplet exterior (red), periphery (blue) and interior (black). Under each experimental condition, fluorescence is highest at the droplet periphery, followed by the droplet interior. The lowest fluorescence intensities are consistently observed exterior to droplets (Fig. 4E). The fluorescence increases nearly linearly with actin concentration both inside and outside the coacervate droplets (Fig. 4E).

In addition to the average partition coefficient (Fig. 1E), we report two additional partition coefficients derived from these intensity profiles; one for the ratio of droplet periphery (maximum observed intensity) to average intensity in the exterior of the droplet (*PC*_Periph_) (Fig. 4F, blue/red), and a second for the ratio of droplet interior intensity (near the center of the droplet) to the average intensity in the exterior (*PC*_Int_) (Fig. 4F, black/red). Both partition coefficients are greater than unity, indicative of partitioning of actin to the polymer-dense coacervate phase from the polymer-dilute phase. *PC*_Periph_ and *PC*_Int_ both tend to reach plateaus for actin concentrations above 0.1 μM; *PC*_Periph_ values grow almost 10-fold before stabilizing at ≈45 once the global actin concentration reached 0.1 μM, while *PC*_Int_ values increase over the range of actin concentrations investigated in the current study, and appear to approach a plateau value of ≈10. The saturation of the *PC*_Periph_ with global actin concentration suggests that exchange of protein between the polymer-dense and dilute phases occurs readily, as has been reported in other liquid phase-separated systems (4).

## DISCUSSION

We present proof-of-concept experiments demonstrating that a polypeptide-based complex coacervate can be used as a model bioreactor to control the localization and activity of the self-assembling cytoskeletal protein actin. We find that actin partitions spontaneously to the coacervate phase, and that its partitioning is not influenced by BSA. Strong partitioning of actin to pLK/pRE coacervates increases the local actin concentration, contributing substantially to a >50-fold increase in the actin assembly rate at the highest concentrations of actin and coacervate. Actin filaments of canonical structure localize to the coacervate periphery, effectively forming core-shell particles, with the actin shell density controlled by the actin concentration.

### Partitioning vs. encapsulation of client proteins

Previous work interpreted the preferential localization of the client protein BSA to the pLK/pRE coacervate phase as “encapsulation” (8, 29, 30). The implication of this language is that exchange of client molecules between the coacervate and dilute phases is either non-existent or so small as to be negligible, as with encapsulation within lipid vesicles or emulsion droplets (31, 32). Indeed, Black et al. argued that entry of the large (66 kDa) client into the coacervate phase requires the formation of an intermediate electrostatic complex between the client and an oppositely-charged polyelectrolyte in solution prior to phase separation, and that client release is triggered by pH-induced dissolution of the coacervate phase (8).

Our present results are more consistent with a molecular view termed partitioning (4), where the partition coefficient reflects the equilibration of steady fluxes of client molecules into and out of the coacervate phase. For instance, we observe partitioning of BSA within ≈30 s upon addition to pre-formed pLK/pRE coacervates (Fig. S3). Given the very low polypeptide concentration in the dilute phase (< 30 nM pLK), this suggests that recruitment of the client to the coacervate does not require the formation of an intermediate complex with a polyelectrolyte. Additionally, the saturation of the partition coefficient for actin concentrations above 0.1 μM (Fig. 4F) is indicative of an equilibration between the client concentrations in the dilute and coacervate phases, which necessarily requires exchange.

Equilibrium partitioning in synthetic polypeptide coacervates is reminiscent of other recent *in vitro* work wherein client proteins of low-valency spontaneously partition into liquid phase-separated structures composed of high-valency scaffold proteins (4), as well as in coacervates formed from natural biopolymers (33). This is particularly interesting in that binding is mediated by specific low-affinity protein-protein interactions in the former case, in contrast to the non-specific electrostatic interactions presumed in the case of coacervates. This suggests that the capacity to selectively partition client molecules may be a general property of condensed liquid-like phases, independent of the interactions driving partitioning.

### Origin of peripheral F-actin localization

Below, we examine three non-mutually exclusive physical mechanisms for the peripheral localization of F-actin in coacervates droplets: filament buckling, macromolecular depletion, and interfacial adsorption.

F-actin does not appear to protrude from micron-sized coacervate droplets, suggesting that coacervate surface tension may play a role in confining F-actin. One mechanism for peripheral filament localization is that surface tension causes filaments to buckle once the contour length exceeds the droplet diameter. A comparison of the energy required to increase the coacervate surface area to accommodate a protruding filament of length L and diameter d with a cylindrical cap, *E_Area_* = *πd* (*L* − *2R*)*γ*, with the energy required to bend the filament into a circular arc with radius R equal to that of the droplet,

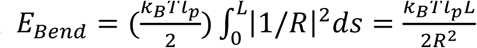, yields the shortest length greater than 2R for which bending is energetically favorable:

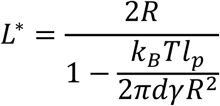

where k_B_ is the Boltzmann constant, T is temperature, and l_p_ = 10 μm is the persistence length of F-actin (34). In a 1-μm diameter coacervate droplet with the surface tension γ = 1 mN/m (17) at room temperature, bending is preferable for filaments longer than ≈1 μm. Coacervate surface tension is thus sufficient to bend F-actin with contour lengths larger than the droplet diameter. However, the observation that filaments and bundles even shorter than the droplet diameter are peripherally localized (Fig 4 A-C, 0.05 μM panels) indicates that surface tension-induced buckling cannot be the sole cause.

The peripheral localization of F-actin is reminiscent of the well-known crowding of F-actin to interfaces observed in the presence of macromolecular crowding agents (35, 36), which arises from depletion interactions (37). To assess this, we estimate the osmotic pressure needed to crowd F-actin to an interface to be Π* ≅ 450 Pa (Supplementary Information). We estimate the osmotic pressure of the coacervate interior as that arising from a solution of flexible polymers characterized by a mesh size (38), ξ, as Π = *k_B_T/ξ*^3^. This suggests that a coacervate with mesh size ξ ≤ 20 nm would generate sufficient osmotic pressure to drive peripheral localization of F-actin. We estimate the mesh size of our pLK/pRE coacervates to be 2-3 nm (Supplementary Information), which supports the plausibility of a depletion-based mechanism. Noting the empirical observation that long filaments crowd more readily than short ones, macromolecular depletion could preferentially crowd long, high aspect ratio filaments and bundles, leaving short filaments, actin monomer, and BSA uniformly distributed.

A third, non-mutually exclusive mechanism for peripheral localization of F-actin is filament adsorption to the coacervate/bulk interface. For example, electrostatic interactions between F-actin and the coacervate could drive adsorption in a process akin to that of polyelectrolyte-mediated emulsion stabilization (39). Alternately, a difference in the interfacial tensions between F-actin and the solution and the coacervate, respectively, could drive localization of filaments to the coacervate/solution interface, such as seen in Pickering emulsions (40). In support of an adhesion-based mechanism, we note that filaments occasionally wrap around coacervate droplets when assembling in solution (Fig. S5, Movie S2), indicative of an attractive interaction between F-actin and the coacervate/solution interface.

Importantly, not all actin fluorescence is peripherally localized. Actin fluorescence intensity in the center of pLK/pRE coacervates is diffuse and enriched by as much as 10-fold compared to the surrounding solution (Fig. 4F). Interior fluorescence increases with global actin concentration, which is inconsistent with the peripheral localization of all filaments. In that case, the interior fluorescence would correspond to solely actin monomers, which we would predict to have a constant local concentration of c_local_ = c* = 0.1 μM at steady-state. This suggests that the coacervate interior contains a mixture of monomers and filaments. Given that a 250-nm filament (~90 actin subunits) cannot be resolved with conventional light microscopy, an interior including both monomers and short filaments is consistent with the diffuse fluorescence signal we observe. The aforementioned mechanisms of peripheral localization all depend on F-actin length. As such, the presence of short filaments in the interior does not qualitatively distinguish between them. Understanding how peripheral localization is regulated will be an exciting avenue for future studies.

### Mechanism of actin assembly enhancement

The assembly of actin within coacervates for global actin concentrations below the critical concentration is largely predicted by the 30-fold increase in the local actin concentration via partitioning into the coacervate phase. Using measured partition coefficients, we calculate the concentrations at which filament assembly within coacervates is expected and find that partitioning is sufficient to explain filament assembly down to 0.05 μM (red dashed line, Fig. 4F). However, our measured partitioning is not quite sufficient to polymerize actin within coacervates at the lowest concentration (0.01 μM), yet we observe strong peripheral intensity (and interpret this to be polymerized actin). It is possible that the coacervate environment may alter the reaction rate kinetics of actin assembly (41, 42), as has been seen for transcription in cell-lysate coacervates (43). Indeed, since the coacervate-phase volume fraction (40%) and viscosity (2 Pa-s) are similar to those of the cytoplasm, this system may serve as a useful platform to study biochemical reactions in a more physiological environment.

### Implications for biochemical reaction regulation

The high local concentrations generated by partitioning provide for an elegant means to both spatially localize and enhance the rates of biochemical reactions. Spontaneous partitioning to a condensed liquid-like phase substantially reduces the quantities of protein needed to study reactions under more physiological conditions. Of particular interest is the possibility of having direct control over partition coefficients and other local physicochemical properties of the coacervate phase as a means to control biochemical reactivity.

In summary, we have illustrated spontaneous partitioning of proteins inside coacervate droplets, leading to markedly increased actin assembly rates and spatial confinement of filaments. Assembly rate enhancements reported here are qualitatively consistent with a model in which these enhancements were contributed by an increase in the local effective concentration of actin monomers in the coacervate phase. Our work introduces exciting avenues for the use of synthetic polymers to control physical and biological properties of bioreactors, and address questions in biology about the biochemistry of molecules in cell-like micro-environments.

## MATERIALS AND METHODS

Solutions containing pLK/pRE coacervates and fluorescently-labeled proteins were imaged on a spinning disk confocal microscope. Details of all experimental methods and analysis can be found in the *SI Materials and Methods*.

## ACKNOWLEDGEMENTS

The authors thank C. Suarez, K. Weirich, T. Witten, and J. Vieregg, as well as the Gardel, Kovar, and Tirrell labs, for helpful discussion and suggestions. We thank L. Li for assistance with gel permeation chromatography measurements. DRK, MLG, and MVT acknowledge support from the University of Chicago MRSEC (NSF DMR-1420709). PMM thanks the University of Chicago MRSEC for a graduate fellowship.

## Supplementary Information

### Polypeptide Synthesis and Storage

The racemic polyanion poly(L,D)-glutamate sodium salt (pRE) was purchased from Alamanda Polymers Inc. This is a 50/50 random copolymer of L- and D-glatamate stereoisomers. Poly(L-lysine hydrochloride) (pLK) was purchased from Alamanda Polymers Inc. or prepared in-house via N-carboxyanhydride (NCA) polymerization and characterized using gel permeation chromatography (GPC) and ^1^H NMR (44). See Table S1 for the source, mean degree of polymerization, molecular weight, polydispersity, for the polypeptides used in each Figure.

Stock solutions were prepared gravimetrically using MilliQ water (resistivity of 18.2 MΩ-cm, Millipore) at concentrations of either 10 or 20 mM with respect to the number of monomers (that is, the number of acid or base groups) present in solution and then adjusted to pH 7.0 using concentrated solutions of HCl and NaOH, as needed. These solutions were stored at 4 °C, and wrapped in parafilm to minimize evaporation. Unless otherwise noted, peptide concentrations are reported in terms of charge equivalents. For instance, a concentration of 5 mM pLK indicates a 5 mM concentration of lysine residues, each with an average charge equivalent of +1 at pH 7. Correspondingly, the concentration of pLK molecules is 50 μM, with each pLK molecule composed of, on average, 100 +1-charged lysine residues.

### Protein Purification, Labeling, and Storage

Actin was purified from rabbit skeletal muscle acetone powder (Pel-Freez) as previously described (45). A subset of gel-filtered actin was labeled on Cys-347 with either Oregon green 488 iodoacetamide (OG), Tetramethylrhodamine-5-maleimide (TMR), or pyrenyl iodoacetamide (Invitrogen) (35, 46, 47). All actins were stored in Calcium Buffer-G (CaBG: 2 mM Tris-HCl, pH 8.0 at 22 °C, 0.2 mM ATP (adenosine triphosphate), 0.5 mM DTT (1,4-Dithiothreitol), 0.1 mM CaCl_2_, 1 mM NaN_3_). Unlabeled, OG- and pyrene-labeled actins were stored at 4 °C, while TMR-labeled actin was flash-frozen with liquid nitrogen and stored at -80 °C. Prior to use, non-frozen actins were dialyzed against 0.2 L fresh CaBG for 18-24 h, and clarified via ultracentrifugation at 177,000 x g (average relative centrifugal force) for 30 minutes at 4 °C. The top 90% of the supernatant was retained, stored on ice, and used within 6 days. TMR-actin was rapidly thawed by hand, and similarly clarified via ultracentrifugation at 177,000 x g for 30 minutes at 4 °C. The top 75% of the supernatant was retained, stored on ice, and used within 2 days.

Bovine Serum Albumin (BSA) (Fraction V, Fisher Scientific) was labeled on exposed cysteines with Alexa Fluor 647 Maleimide (Invitrogen) according to the manufacturer’s protocol, and stored in 1x Phosphate-Buffered Saline (PBS, pH 7.5 at room temperature) at -80 °C. Prior to use, labeled BSA was thawed rapidly by hand, and clarified by centrifugation at 21,000 x g for 20 minutes at 4 °C. The entire supernatant was retained, stored at 4 °C, and used within one month.

### Protein Concentration Determination

Protein concentrations were determined spectrophotometrically. Absorbance measurements were made using an Ultrospec 2100 pro (Amersham Biosciences) or a NanoDrop ND-1000 (Thermo Scientific) UV/VIS Spectrophotometer with optical path lengths b = 1 cm and b = 0.1 cm, respectively. Sample absorbance was converted to protein concentration (and corrected for label absorbance, as appropriate) using the following expressions:

[Unlabeled-actin] = <*A_290_*> *(38.5 μM cm) / b (48)

For OG-labeled actin (35):

[Total actin] = (<*A_290_*> - (0.17*<*A_491_*>))*(38.5 μM cm) / b
[OG-actin] = <*A_491_*> / ((0.0778 μM^-1^ cm^-1^)*b)

For TMR-labeled actin:

[Total actin] = (<*A_290_*> - (0.1185* <*A_557_*>))*(38.5 μM cm) / b
[TMR-actin] = <*A_557_*> / ((0.1009 μM^-1^ cm^-1^)*b)

For Pyrene-labeled actin (48):

[Total actin] = (<*A_290_*> - (0.127*<*A_344_*>))*(38.5 μM cm) / b
[Pyrene-actin] = <*A_344_*>*(45.0 μM cm) / b

For BSA, the molar extinction coefficient was estimated to be *E_280_* = 40,800 M^-1^ cm^-1^ using the online tool ProtParam (http://web.expasy.org/protparam/), the amino acid sequence associated with the UniProt ID P02769 for *Bos taurus* serum albumin, and assuming reduced Cys residues. The approximate extinction coefficient for Alexa Fluor 647 C_2_ maleimide is *E_650_* = 265,000 M^-1^ cm^-1^ (49). The concentration of BSA and Alexa Fluor 647 were thus calculated according to:

[Total BSA] = (<*A_280_*> -(0.03*<*A_651_*>)) / ((0.0408 μM^-1^ cm^-1^)*b)
[Alexa 647] = <*A_651_*> / ((0.265 μM^-1^ cm^-1^)*b)

### Sample Preparation

Mg-ATP-actin is prepared in Tube A from Ca-ATP-actin by 2-minute incubation at room temperature (RT) with 1/10th volume 10x Magnesium Exchange buffer (1x ME buffer: 50 μM MgCl_2_, 200 μM EGTA (ethylene glycol-bis(beta-aminoethyl ether)-N,N,N′,N′-tetraacetic acid) in MilliQ-purified water, stored at RT). Mg-ATP-actin is then incubated at RT with either Latrunculin A dissolved in DMSO (dimethyl sulfoxide) for 5 min or an equivalent volume of DMSO, as a blank, for minimal time (< 30 s), followed by addition of pLK (or a blank of MilliQ-purified water) and incubation for 2-3 min at RT. Tube B is assembled from 10x KMEI buffer (1x KMEI: 50 mM KCl, 1 mM MgCl_2_, 1 mM EGTA, 10 mM Imidazole, pH 7.0, stored at RT), Magnesium Buffer-G (MgBG: 2 mM Tris-HCl, pH 8.0, 22 °C, 0.2 mM ATP, 0.5 mM DTT, 0.1 mM MgCl_2,_ 1 mM NaN_3_, stored on ice), and pRE (or a blank of MilliQ-purified water). Polyelectrolyte phase separation and actin polymerization is then initiated simultaneously through the addition of Tube B to Tube A and rapid mixing by pipette. Samples with ~ 1 mM or more total polypeptide are visibly turbid < 2 s after addition of Tube B to Tube A.

### Microscopy

Samples were loaded into flow cells constructed from mPEG-Silane-treated glass and double-sided tape, as described previously (50). Samples were imaged at room temperature (~24 °C) on an inverted microscope (Ti-Eclipse; Nikon, Melville, NY) equipped with a confocal scan head (CSU-X, Yokogawa Electric, Musashino, Tokyo, Japan), a laser merge module (LMM5, Spectral Applied Research, Richmond Hill, Ontario, Canada) containing 488-nm, 560-nm, and 635-nm laser lines for fluorescence imaging, as well as polarizers, Nemarski prisms, and a white transmitted-light source for differential interference contrast (DIC) imaging. Images were formed using a 60x, DIC-compatible, water-immersion objective (Nikon, Melville, NY) with a numerical aperature (NA) of 1.2, and corrections for apochromatic and flat field aberrations (Plan Apo). Images were acquired on a scientific complementary metal-oxide-semiconductor (sCMOS) camera (Zyla 4.2; Andor Technologies, Belfast, Northern Ireland), with a physical pixel size of 6.5 microns per side. All imaging hardware was controlled using METAMORPH acquisition software (Molecular Devices, Eugene, OR).

### Image Analysis

All quantitative image analysis was performed using imageJ (NIH, imagej.nih.gov/ij/) and custom code written in MATLAB (MathWorks, Natick, MA). Camera dark noise (100 counts) was subtracted from all fluorescence images prior to analysis.

#### Measurement of average partition coefficient

Following camera noise subtraction, fluorescence images were corrected for uneven illumination (51) using a reference fluorescence image of a plastic slide. Two masks were created for each corrected fluorescence image by thresholding. The first is called the background mask, and the second is called the droplet mask. Note that in order to reduce the impact of imaging artifacts which result from spinning-disk confocal imaging, such as pin-hole crosstalk, the background mask is not simply the inverse of the droplet mask (4, 52). We used the modal pixel intensity from the illumination-corrected fluorescence image as the threshold for the background mask. After application of the threshold, the close, invert, close, and fill holes operations were applied in ImageJ. This procedure yields a mask which excludes droplets, as well as regions outside of droplets where systematic pin-hole cross-talk effects are visible.

To create the droplet mask, the illumination-corrected image was thresholded again, this time using the average full-width-at-half-max intensity obtained from linescans across 5-10 coacervate droplets in the field of view. After application of the threshold, the invert, close, fill holes, and invert operations were applied in ImageJ.

Individual candidate droplets were then automatically identified from the droplet mask using custom code written in MATLAB. Candidate droplets were filtered by size and eccentricity. As a result of the strong peripheral localization of actin fluorescence, candidate droplets occasionally had a crescent-like morphology, similar to an unclosed circle. This occurred if the fluorescence at some position along the droplet boundary was below the masking threshold. To remove droplets for which the mask gave unclosed circles, candidate droplets in actin fluorescence images were further filtered by removing candidates for which the center of mass fell outside the candidate droplet region. The size and average intensity of each filtered droplet was then computed using the droplet mask and the illumination-corrected fluorescence image. The average background intensity was computed using the background mask and the illumination-corrected image. Finally, the average partition coefficient for each droplet was calculated as the ratio of the average intensity within the droplet mask to the average background intensity.

#### Measurement of peripheral and interior partition coefficients

Fluorescence intensity linescans are calculated using a transverse width of 5 pixels along paths indicated in the accompanying image, and normalized according to
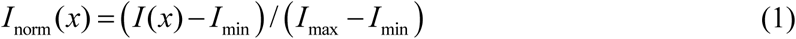

where *I(x)* is the intensity at position *x* along the path, and *I*_min_ and *I*_max_ are the minimum and maximum values of the intensity along the path, respectively. Interior, Periphery, and Exterior fluorescence values (e.g. Fig. 4F) are obtained from individual linescans. Interior is the average of a 5-pixel segment along the linescan, co-centered with the droplet. Periphery is the average of the maximum intensity along the linescan from either side of the droplet center. Exterior signal is calculated on each side of the droplet as the average intensity along the 6-pixel segment of the linescan extending from 10 pixels beyond the periphery peak to 15 pixels beyond the peak. The reported Exterior value is the average of the Exterior signals from each side of the linescan. One-sided Exterior values (as opposed to the average of both sides) are reported when the relevant segment on one side of the linescan impinges upon an adjacent droplet. Linescan orientation is chosen to minimize the frequency of this. Partitioning coefficients (4) are calculated as the ratio of either Periphery or Interior fluorescence to Exterior fluorescence.

#### Measurement of fluorescence intensity correlation coefficients

Calculation of fluorescence intensity correlation coefficients is restricted to include only pixels within droplets using a mask for each field of view. Masks were generated in ImageJ from DIC images taken at the approximate midplane of the droplet of interest by application of a Gaussian blur (radius 1.00), threshold-based conversion to a binary image, and the sequential application of the binary operations close, fill holes, and dilate, where dilate is applied twice. The intensities in the fluorescence cross-correlation histogram (e.g. Fig. 2C) are normalized independently in each channel according to equation (1) above, where the domain of *x* is all pixels identified from the mask as within droplets.

### Fluorescence Spectroscopy

A stock solution of 15 μM Ca-ATP-actin (typically 10-20 % pyrene-labeled) is prepared from solutions of unlabeled and highly-labeled (typically > 90 %) actin in a 1.5-mL microcentrifuge tube, and then converted from Ca-ATP-actin by 2-minute incubation with 1/10th volume 10x ME buffer and 1/10th volume of 100x anti-foam (Antifoam 204, Sigma) at RT. Mg-ATP-actin is then incubated with the desired concentration of pLK (or a blank of MilliQ-purified water) for 2- 3 min at RT in up to 12 rows of a 96-well plate (Assay Plate 3686, Corning). A multichannel pipette is then used to simultaneously deliver solution assembled from 10x KMEI buffer MgBG, and pRE (or a blank of MilliQ-purified water) to the wells containing actin, thereby initiating assembly (through the addition of MgCl_2_ and KCl) and phase separation (through the addition of the polyanion) simultaneously. The final reaction volume is 150 μL. Actin assembly is monitored at RT by pyrene fluorescence (Excitation: 365 nm, Emission: 407 nm) in a fluorescence plate reader (Saphire2, Tecan). The timecourse of pyrene excimer fluorescence (Excitation: 343 nm, Emission: 478 nm (24) was monitored concomitantly with assembly by sequentially alternating the excitation/emission wavelength pairs on a fluorescence plate reader (Infiniti m200, Tecan). We report the reciprocal of the time to 50 %-assembly (1/t_1/2_) as a measure of the mean assembly rate. t_1/2_ is determined for each 407-nm fluorescence timecourse as the time-point at which the fluorescence intensity is closest to *I*_1/2_, which is defined as

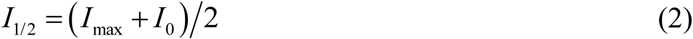

where *I*_max_ is the maximum intensity of the timecourse, and *I*_0_ is the initial fluorescence value of the actin-only control sample, which is an estimate of the sum of background contributions from the detector, buffers, and pyrene-actin monomers for all conditions.

### Physical Estimates

#### Estimation of osmotic pressure required to crowd F-actin to an interface

We estimate the minimum osmotic pressure needed to crowd F-actin to an interface as that of a 0.25 % (w/v) methylcellulose solution, which is known to be sufficient (36). Using the van ‘t Hoff formula Π = *cN_A_k_B_T* for the osmotic pressure of a dilute polymer at molar concentration *c*, where N_A_ is Avagadro’s number, we find that a concentration of *c* = 0.25% ≅ 180 *μM* of 14-kDa methylcellulose gives a pressure of Π* ≅ 450 Pa at room temperature.

#### Estimation of polypeptide species concentration in dilute phase

The dilute phase concentration of a polypeptide is the critical concentration for phase separation at that temperature and salt concentration. Thus, observation of droplet formation upon mixing of pLK with pRE, each at a final global concentration X, indicates that X is greater than the critical concentration, and thereby greater than the dilute phase concentration. pLK/pRE coacervate droplets are visible by DIC at concentrations of 30 μM per polypeptide and above. However, punctate BSA fluorescence is observed at 3 μM per polypeptide, even though droplets are not readily discernible by DIC. This suggests that phase separation does still occur at this concentration. We thus use 3000 nM / 100 residues per polypeptide = 30 nM as a conservative upper bound on the dilute phase concentration for each polypeptide chain species.

#### Estimation of polypeptide concentration in coacervate phase

From dimensional analysis, we estimate the polypeptide concentration in the coacervate phase from the mass fraction according to *c* = *ρ_pp_f_pp_ / M_W_*, where *ρ_pp_* is the mass density of a solid polypeptide substance, *f_pp_* is the polypeptide mass fraction in the coacervate phase, and *M_W_* is the polypeptide molecular weight. We estimate the polypeptide mass density *ρ_pp_* the density of solid glutamic acid, *ρ_E_* = 1538 mg/mL (53). Unpublished measurements of *f_pp_* for pLK/pRE coacervates under similar conditions find a total polypeptide mass fraction of 0.38, or 0.19 per polypeptide species. This is comparable to previously published reports on other coacervate systems, where the water content was reported to saturate at ~ 60 % (54). Using these values, and approximating the molecular weight as 12.9 kDa (Table S1), yields a polypeptide chain concentration of ~2 mM per species, corresponding to a peptide concentration of ~ 2 M per species. In the main text, we thus report the pLK concentration estimate as being on the order of 1-3 M in the coacervate phase.

#### Estimation of coacervate mesh size

We estimate the mesh size as (38) 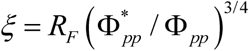, where *R_F_* is the root-mean-square end-to-end distance of the polypeptide, 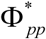 is the polypeptide volume fraction at which spheres of radius *R_F_* containing one polymer each begin to overlap, and is the polypeptide volume fraction of the coacervate phase. Treating the polypeptides as self-avoiding flexible chains in good solvent, we have that (38) *R_F_* = *aN^v^* = *aN*^3/5^, where *a* is the effective monomer size of the self-avoiding chain, N is the number of such monomers per chain, and *v* is the Flory exponent. We also have that 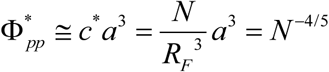, where *c*\* is the overlap polymer concentration. Combining these expressions gives (38) *ξ* = *a*Φ*_pp_*^−3/4^.

We take the effective monomer size of a polypeptide in the coacervate phase to be the Kuhn length, which we estimate as (55) 0.8 +/- 0.1 nm. We estimate the volume fraction as
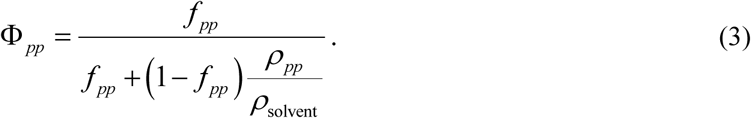

Taking the total polypeptide mass fraction to be *f_pp_* = 0.38 as above, and approximating *ρ*_solvent_ as 1000 mg/mL, we estimate the volume fraction as Φ*_pp_* = 0.28. Combined with the Kuhn length estimate, this gives a mesh size estimate of ~2.1 nm. In the main text, we estimate the mesh-size as 2-3 nm in the coacervate phase.

**Table S1.** Summary of polypeptides used in this study.

| **Polypeptide** | **Source** | **Degree of Polymerization** | **Molecular Weight (kDa)** | **Polydispersity** | **Experiments** |
| --- | --- | --- | --- | --- | --- |
| pLK100 | purchased | 100 | 12.8 | 1.004 | Fig. 1, 2, 3D, 4<br>Fig. S1, S2, S3, S4 |
| pLK126 | in-house | 126 | 16.2 | 1.30 | Fig. 3B-C, S5 |
| pRE100 | purchased | 100 | 12.9 | 1.15 | Fig. 1, 2, 3D, 4<br>Fig. S1, S2, S3, S4 |
| pRE98 | purchased | 98 | 12.7 | 1.02 | Fig. 3B-C, S5 |

**Figure. S1.**
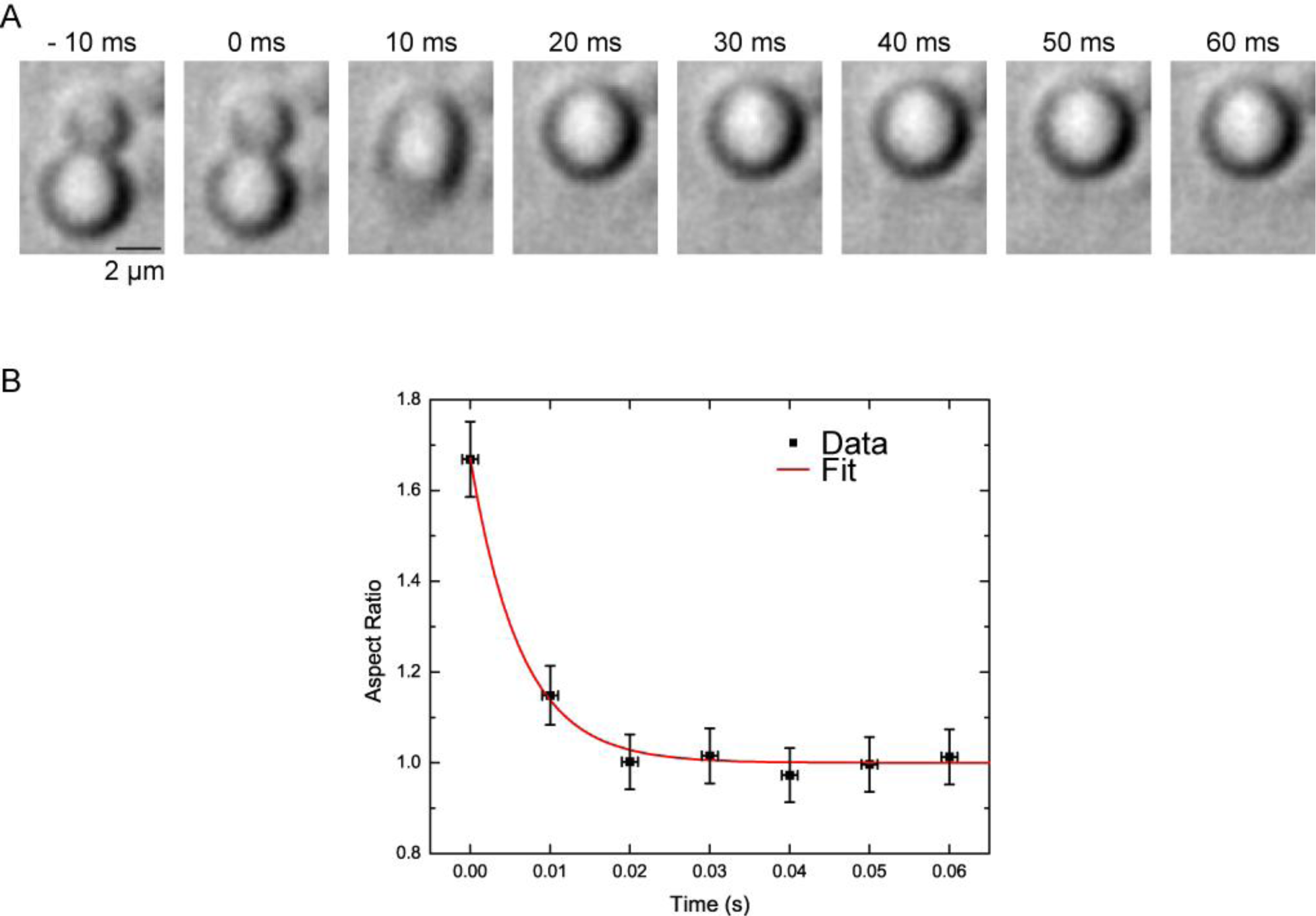
Liquid-like properties of pLK/pRE coacervates. (A) DIC microscopy timelapse of two coacervates merging. Both droplets are initially round, suggestive of surface-tension dominated shapes. Upon fusing, the initially dumbbell-shaped interface rapidly relaxes to a round, surface-area-minimizing conformation of diameter d = 4 μm. (B) Aspect ratio of droplets in (A) during coalescence (black). Red line is an exponential fit, with a time constant of τ = 6.33 ms. We estimate the inverse capillary velocity as *v*^−1^ = *τ/d* = 1.6 ms/μm. Error bars represent uncertainty in aspect ratio from a single coalescence event (dy) and acquisition time (dx). Conditions are 5 mM pLK, 5 mM pRE, 50 mM KCl, 1 mM MgCl_2_, 1 mM EGTA, 10 mM imidazole (pH 7.0), and 80 μM ATP (all concentrations final).

**Figure. S2.**
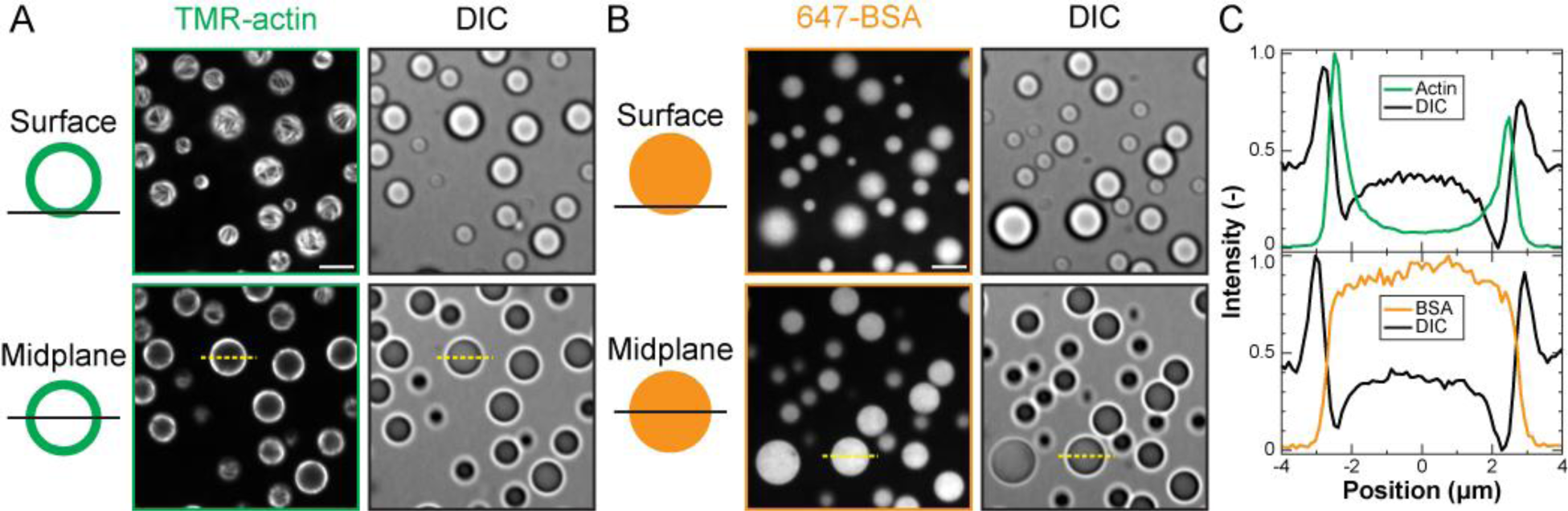
Peripheral localization of F-actin in polypeptide coacervates is BSA-independent. (A-B) Confocal fluorescence (left) and DIC (right) micrographs of polypeptide coacervates containing either TMR-actin (A, red) or 647-BSA (B, cyan) on non-adherent substrates. Top row is focused at the interface of the coacervates and the substrate (surface), and bottom row is approximately the droplet midplane. Scale bar is 5 μm. (C) Normalized fluorescence intensity linescans along the dashed yellow lines indicated in (A) (top) and in (B) (bottom). Conditions are 0.5 μM protein (either Mg-ATP-actin (47% TMR-labeled) or BSA (91% Alexa-647-labeled)) incubated with 5 mM pLK prior to addition of 5 mM pRE in 50 mM KCl, 1 mM MgCl_2,_ 1 mM EGTA, 10 mM imidazole (pH 7.0), and 72 μM ATP (all concentrations final). Note that the average partition coefficients for (A) and (B) are reported in Figure 1 of the main text.

**Figure S3.**
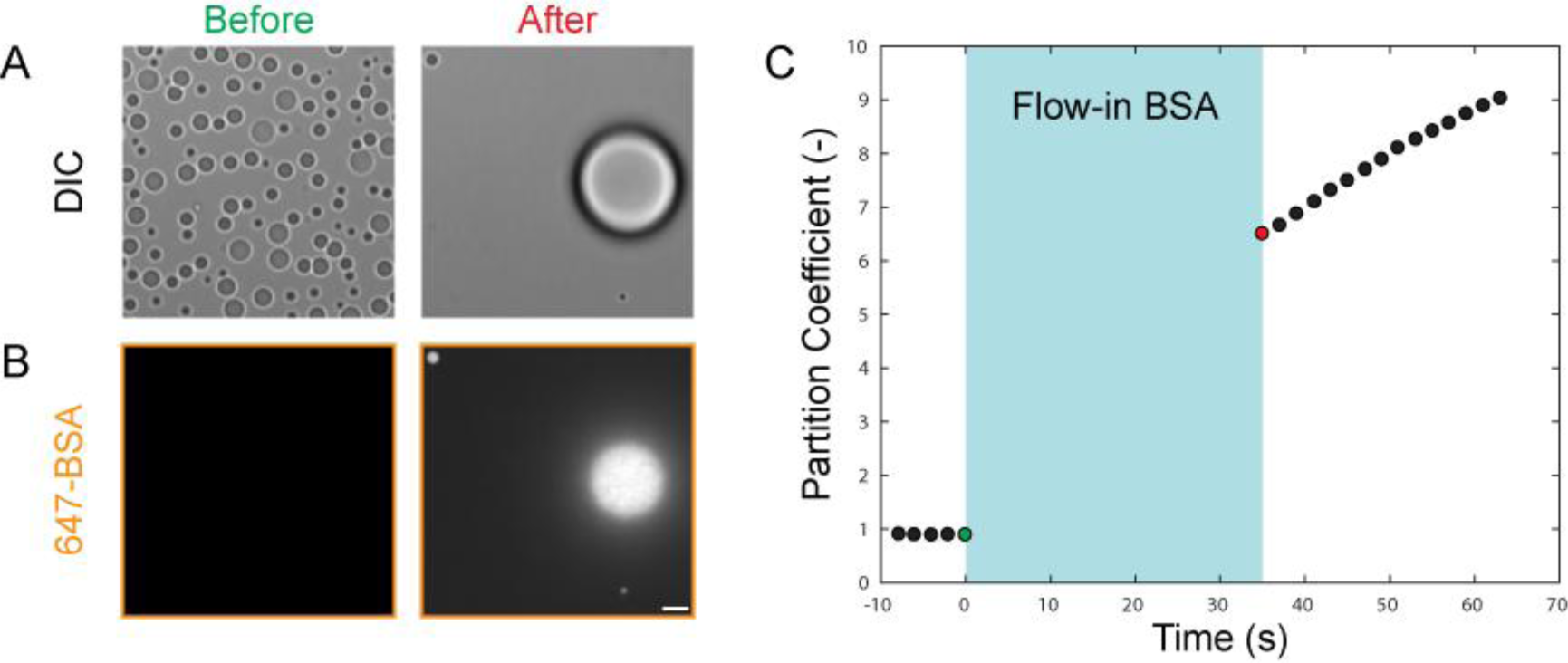
Flow-in of BSA to pre-formed pLK/pRE coacervates. (A-B) Time-lapse confocal DIC (A) and fluorescence (B) micrographs of pLK/pRE coacervates before (left, green) and immediately after (right, red) the addition of 1.5 μM 647-BSA to the flow-cell chamber. Flow causes many of the sedimented droplets to fuse into larger droplets (A). While only background fluorescence is detectable prior to addition of BSA (B, left), BSA is partitioned to and uniformly localized within the few coacervate droplets remaining in the field of view within 35 s. Scale bar is 5 μm. (C) The partition coefficient of the large droplet visible after BSA flow-in increases with time, but is already above 6 in the first image acquired. Conditions are 5 mM pLK, 5 mM pRE, 50 mM KCl, 1 mM MgCl_2_, 1 mM EGTA, 10 mM imidazole (pH 7.0), and 72 μM ATP (all concentrations final). 1.5 μM 647-BSA is perfused into the chamber in buffer containing 50 mM KCl, 1 mM MgCl_2_, 1 mM EGTA, 10 mM imidazole (pH 7.0), and 72 μM ATP (all concentrations final).

**Figure S4.**
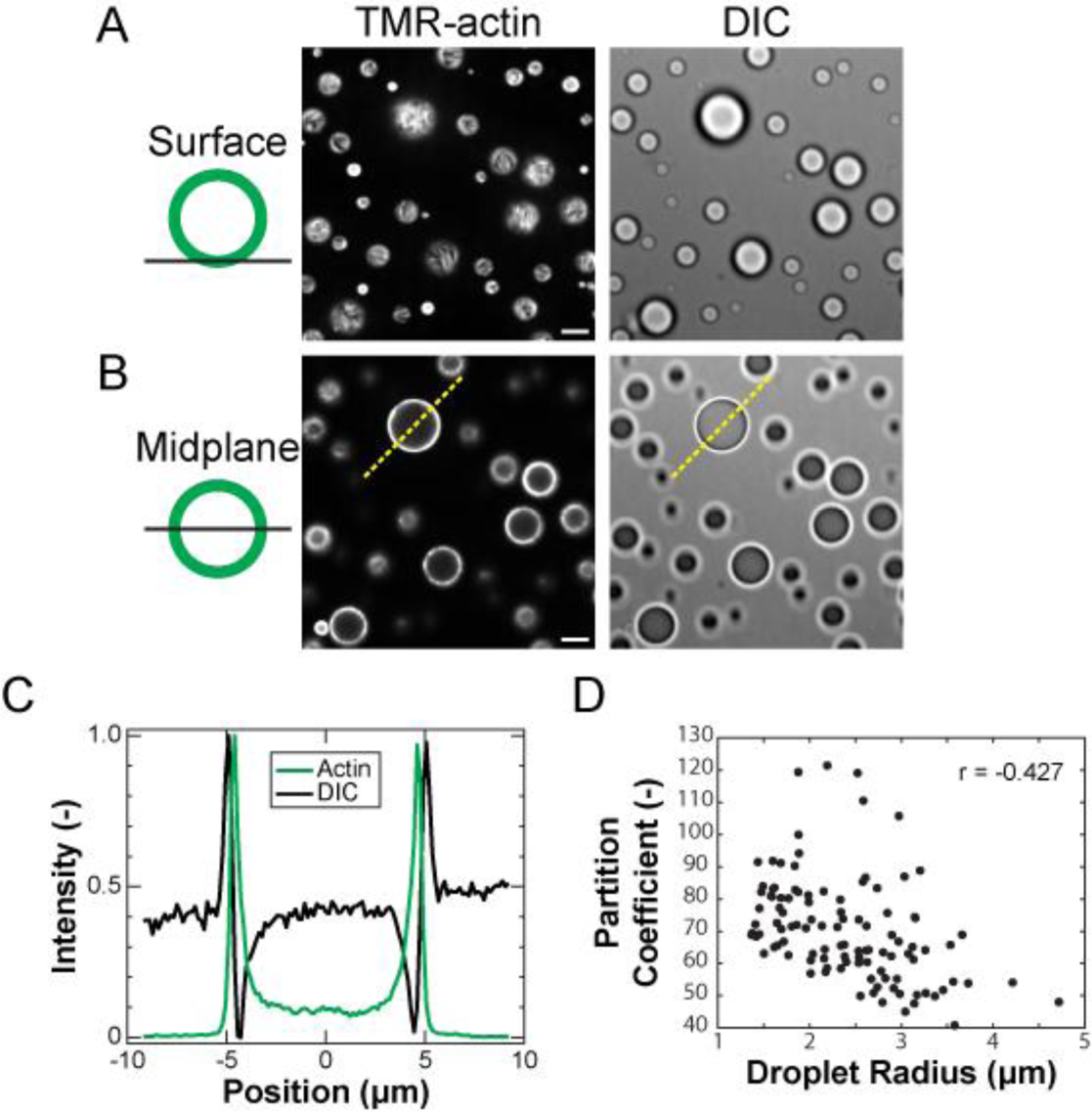
Partitioning and localization patterns are robust to order of addition. (A-B) Confocal fluorescence (left) and DIC (right) micrographs of polypeptide coacervates containing TMR-actin (green) on non-adherent substrates. (A) is focused at the interface of the coacervates and the substrate (surface), and (B) is approximately the droplet midplane. Scale bar is 5 μm. (C) Normalized fluorescence intensity linescans along the dashed yellow lines indicated in (B). (D) Partition coefficients as a function of droplet radius, calculated for a total of N = 117 individual pLK/pRE coacervates with TMR-actin added immediately following phase separation. The average partitioning coefficient is 70.3 +/- 15.9. Error bars denote the standard error of the mean. The Pearson’s correlation coefficient for the data in (D) is r = -0.427, indicating a lower average partition coefficient for larger droplets. Conditions are 5 mM pLK, 5 mM pRE, 50 mM KCl, 1 mM MgCl_2_, 1 mM EGTA, 10 mM imidazole (pH 7.0), and 72 μM ATP (all concentrations final). Following mixing and subsequent phase separation, this solution is added to a small volume of Mg-ATP-actin (47% TMR-labeled), and imaged. The final (global) concentration of actin is 0.5 μM.

**Figure S5.**
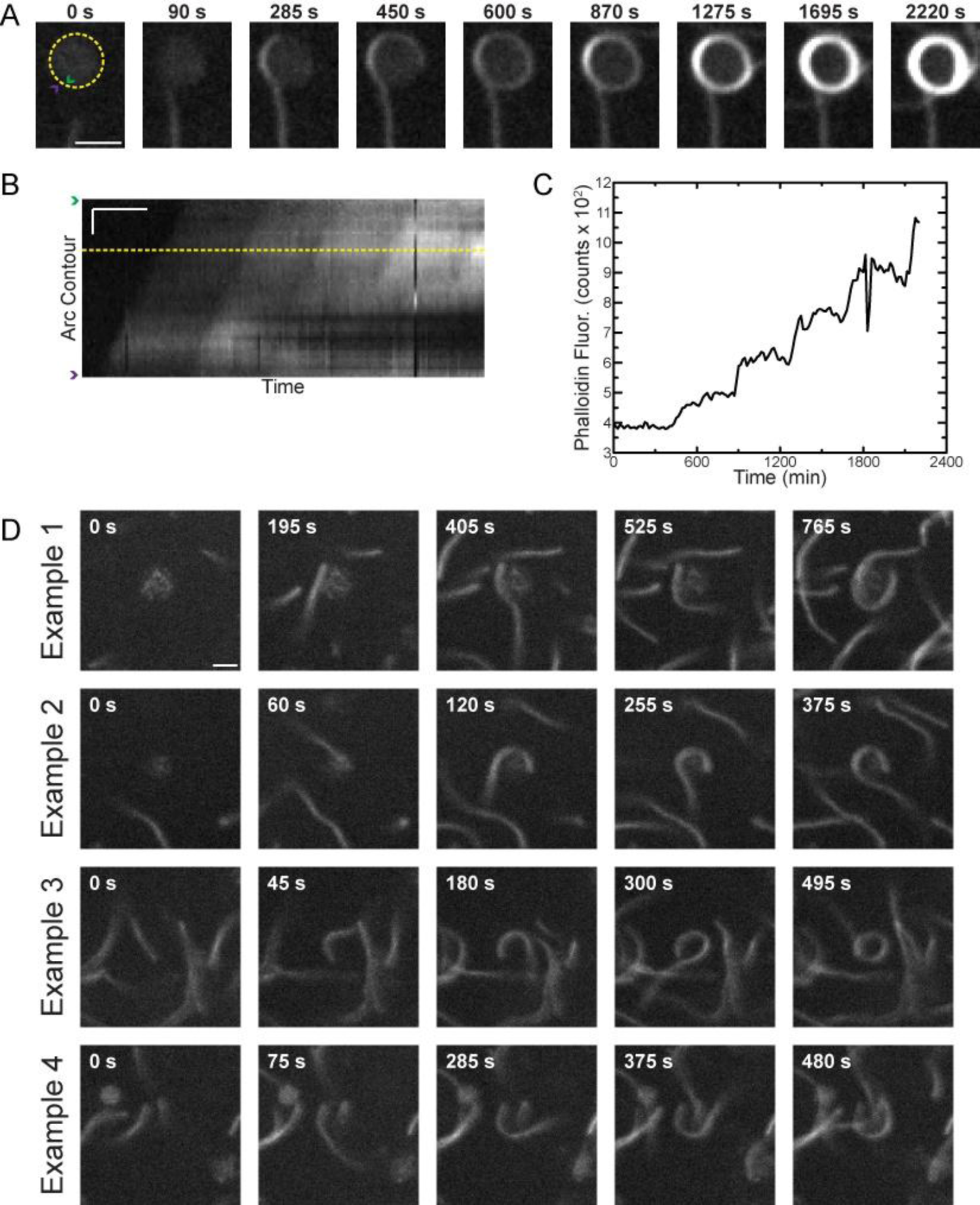
Adhesion of soluble F-actin to the coacervate interface. (A-D) 0.3 μM Mg-ATP-actin (unlabeled) added to a solution of 6 mM pLK/pRE coacervates in 50 mM KCl, 1 mM MgCl_2_, 1 mM EGTA, 10 mM imidazole (pH 7.0), and 72 μM ATP (all concentrations final). (A-C) Also contain 0.5 % (w/v) 14 kDa methylcellulose (MC) and oxygen scavenging system. (A) Fluorescence time-lapse of a growing actin filament approaching a presumed coacervate droplet in presence of MC. Filament contacts coacervate, and winds around the coacervate 4 times upon further elongation. (B) Kymograph along circular yellow path in (A). (C) Intensity along yellow linescan in (B). Step-like increases indicate successive windings. (D) Examples of winding in the absence of MC. Scale bars: 2 μm (A); 1 μm and 300 s (B), 2 μm (D).

**Movie S1.**
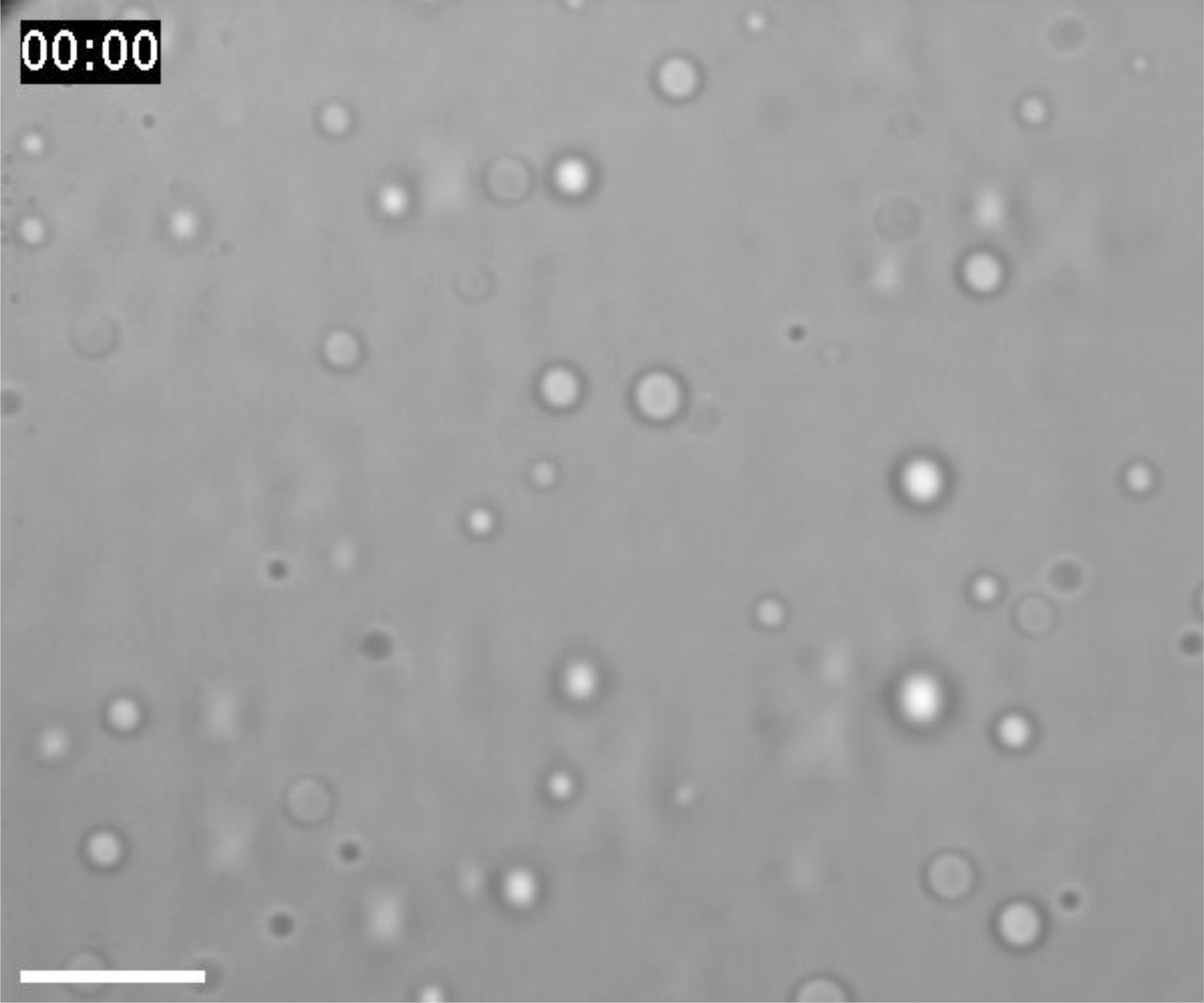
Coacervate droplet coalescence. Scale bar is 10 μm. Playback is 30 frames per second. Time stamp format is mm:ss. DIC timelapse imaging of pLK/pRE coacervate droplets in solution sedimenting onto a passivated glass coverslip due to gravity. Many droplets coalescence, merging into larger droplets. Conditions are 5 mM pLK, 5 mM pRE in 50 mM KCl, 1 mM MgCl_2_, 1 mM EGTA, 10 mM imidazole (pH 7.0), and 72 μM ATP (all concentrations final).

**Movie S2.**
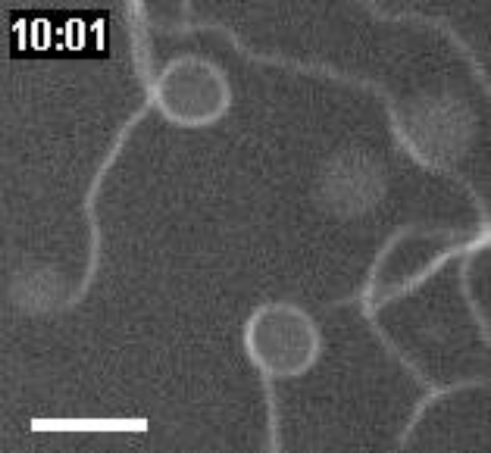
Adhesion of soluble F-actin to the coacervate interface. Scale bar is 4 μm. Playback is 30 frames per second. Time stamp format is mm:ss. Fluorescence time-lapse imaging of growing actin filaments winding around presumed coacervate droplets in the presence of methylcellulose (MC). An image montage of the droplet in the lower center of the field of view is presented in Fig. S5 A. Conditions are 0.3 μM Mg-ATP-actin (unlabeled) added to a solution of 6 mM pLK/pRE coacervates in 50 mM KCl, 1 mM MgCl_2_, 1 mM EGTA, 10 mM imidazole (pH 7.0), 72 μM ATP, 0.5 % (w/v) 14 kDa methylcellulose (MC) and an oxygen scavenging system (all concentrations final).

## REFERENCES

1. Alberts B, et al. (2007) Molecular Biology of the Cell (Garland Science, New York, NY). 5th Ed.

2. Brangwynne CP, et al. (2009) Germline P Granules Are Liquid Droplets That Localize by Controlled Dissolution/Condensation. Science 324(5935):1729–1732.

3. Mitrea DM, Kriwacki RW (2016) Phase separation in biology; functional organization of a higher order. Cell Commun Signal 14:1.

4. Banani SF, et al. (2016) Compositional Control of Phase-Separated Cellular Bodies. Cell 166(3):651–663.

5. Feric M, et al. (2016) Coexisting Liquid Phases Underlie Nucleolar Subcompartments. Cell 165(7):1686–1697.

6. Gucht J van der, Spruijt E, Lemmers M, Cohen Stuart MA (2011) Polyelectrolyte complexes: Bulk phases and colloidal systems. J Colloid Interface Sci 361(2):407–422.

7. Srivastava S, Tirrell MV (2016) Polyelectrolyte Complexation. Advances in Chemical Physics, eds Rice SA, Dinner AR (John Wiley & Sons, Inc., Hoboken, NJ), pp 499–544.

8. Black KA, et al. (2014) Protein Encapsulation via Polypeptide Complex Coacervation. ACS Macro Lett 3(10):1088–1091.

9. de Kruif CG, Weinbreck F, de Vries R (2004) Complex coacervation of proteins and anionic polysaccharides. Curr Opin Colloid Interface Sci 9(5):340–349.

10. Lindhoud S, Claessens MMAE (2015) Accumulation of small protein molecules in a macroscopic complex coacervate. Soft Matter 12(2):408–413.

11. Liu Y, Winter HH, Perry SL (2017) Linear viscoelasticity of complex coacervates. Adv Colloid Interface Sci 239:46–60.

12. Aumiller Jr WM, Keating CD (2016) Phosphorylation-mediated RNA/peptide complex coacervation as a model for intracellular liquid organelles. Nat Chem 8(2):129–137.

13. Koga S, Williams DS, Perriman AW, Mann S (2011) Peptide–nucleotide microdroplets as a step towards a membrane-free protocell model. Nat Chem 3(9):720–724.

14. Pacalin NM, Leon L, Tirrell M (2016) Directing the phase behavior of polyelectrolyte complexes using chiral patterned peptides. Eur Phys J Spec Top 225(8–9):1805–1815.

15. Perry SL, et al. (2015) Chirality-selected phase behaviour in ionic polypeptide complexes. Nat Commun 6:6052.

16. Ou Z, Muthukumar M (2006) Entropy and enthalpy of polyelectrolyte complexation: Langevin dynamics simulations. J Chem Phys 124(15):154902.

17. Priftis D, Farina R, Tirrell M (2012) Interfacial Energy of Polypeptide Complex Coacervates Measured via Capillary Adhesion. Langmuir 28(23):8721–8729.

18. Brangwynne CP, Mitchison TJ, Hyman AA (2011) Active liquid-like behavior of nucleoli determines their size and shape in Xenopus laevis oocytes. Proc Natl Acad Sci 108(11):4334–4339.

19. ProtParam E ExPASy - ProtParam tool. Available at: http://web.expasy.org/protparam/ [Accessed April 21, 2017].

20. Vandekerckhove J, Deboben A, Nassal M, Wieland T (1985) The phalloidin binding site of F-actin. EMBO J 4(11):2815–2818.

21. Tang JX, Janmey PA (1996) The Polyelectrolyte Nature of F-actin and the Mechanism of Actin Bundle Formation. J Biol Chem 271(15):8556–8563.

22. Cooper JA, Walker SB, Pollard TD (1983) Pyrene actin: documentation of the validity of a sensitive assay for actin polymerization. J Muscle Res Cell Motil 4(2):253–262.

23. Sept D, McCammon JA (2001) Thermodynamics and Kinetics of Actin Filament Nucleation. Biophys J 81(2):667–674.

24. Bubb MR, et al. (2002) Polylysine Induces an Antiparallel Actin Dimer That Nucleates Filament Assembly CRYSTAL STRUCTURE AT 3.5-Å RESOLUTION. J Biol Chem 277(23):20999–21006.

25. Brown SS, Spudich JA (1979) Nucleation of polar actin filament assembly by a positively charged surface. J Cell Biol 80(2):499–504.

26. Oriol-Audit C (1978) Polyamine-Induced Actin Polymerization. Eur J Biochem 87(2):371–376.

27. Fujiwara I, Vavylonis D, Pollard TD (2007) Polymerization kinetics of ADP- and ADP-Pi-actin determined by fluorescence microscopy. Proc Natl Acad Sci 104(21):8827–8832.

28. Pollard TD (1986) Rate constants for the reactions of ATP- and ADP-actin with the ends of actin filaments. J Cell Biol 103(6):2747–2754.

29. Nolles A, et al. (2015) Encapsulation of GFP in Complex Coacervate Core Micelles. Biomacromolecules 16(5):1542–1549.

30. Obermeyer AC, Mills CE, Dong X-H, Flores RJ, Olsen BD (2016) Complex coacervation of supercharged proteins with polyelectrolytes. Soft Matter 12(15):3570–3581.

31. Vieregg JR, Tang T-YD (2016) Polynucleotides in cellular mimics: Coacervates and lipid vesicles. Curr Opin Colloid Interface Sci 26:50–57.

32. Kakran M, Antipina MN (2014) Emulsion-based techniques for encapsulation in biomedicine, food and personal care. Curr Opin Pharmacol 18:47–55.

33. Pak CW, et al. (2016) Sequence Determinants of Intracellular Phase Separation by Complex Coacervation of a Disordered Protein. Mol Cell 63(1):72–85.

34. McCullough BR, et al. (2011) Cofilin-Linked Changes in Actin Filament Flexibility Promote Severing. Biophys J 101(1):151–159.

35. Kuhn JR, Pollard TD (2005) Real-Time Measurements of Actin Filament Polymerization by Total Internal Reflection Fluorescence Microscopy. Biophys J 88(2):1387–1402.

36. Murrell MP, Gardel ML (2012) F-actin buckling coordinates contractility and severing in a biomimetic actomyosin cortex. Proc Natl Acad Sci 109(51):20820–20825.

37. Asakura S, Oosawa F (1954) On Interaction between Two Bodies Immersed in a Solution of Macromolecules. J Chem Phys 22(7):1255–1256.

38. de Gennes P-G (1979) Scaling Concepts in Polymer Physics (Cornell University Press, Ithaca, NY). 1st Ed.

39. Andelman D, Joanny J-F (2000) Polyelectrolyte adsorption. Comptes Rendus Académie Sci - Ser IV - Phys 1(9):1153–1162.

40. Bon SAF (2014) CHAPTER 1:The Phenomenon of Pickering Stabilization: A Basic Introduction. Particle-Stabilized Emulsions and Colloids, pp 1–7.

41. Drenckhahn D, Pollard TD (1986) Elongation of actin filaments is a diffusion-limited reaction at the barbed end and is accelerated by inert macromolecules. J Biol Chem 261(27):12754–12758.

42. Frederick KB, Sept D, De La Cruz EM (2008) Effects of Solution Crowding on Actin Polymerization Reveal the Energetic Basis for Nucleotide-Dependent Filament Stability. J Mol Biol 378(3):540–550.

43. Sokolova E, et al. (2013) Enhanced transcription rates in membrane-free protocells formed by coacervation of cell lysate. Proc Natl Acad Sci 110(29):11692–11697.

44. Kramer JR, Deming TJ (2010) General Method for Purification of α-Amino acid-N-carboxyanhydrides Using Flash Chromatography. Biomacromolecules 11(12):3668–3672.

45. Spudich JA, Watt S (1971) The Regulation of Rabbit Skeletal Muscle Contraction I. BIOCHEMICAL STUDIES OF THE INTERACTION OF THE TROPOMYOSINTROPONIN COMPLEX WITH ACTIN AND THE PROTEOLYTIC FRAGMENTS OF MYOSIN. J Biol Chem 246(15):4866–4871.

46. Kovar DR, Kuhn JR, Tichy AL, Pollard TD (2003) The fission yeast cytokinesis formin Cdc12p is a barbed end actin filament capping protein gated by profilin. J Cell Biol 161(5):875–887.

47. Kudryashov DS, Reisler E (2003) Solution Properties of Tetramethylrhodamine-Modified G-Actin. Biophys J 85(4):2466–2475.

48. Hansen S, Zuchero JB, Mullins RD (2013) Cytoplasmic Actin: Purification and Single Molecule Assembly Assays. Adhesion Protein Protocols, Methods in Molecular Biology., ed Coutts AS (Humana Press), pp 145–170.

49. Molecular Probes The Molecular Probes Handbook. Available at: https://www.thermofisher.com/us/en/home/references/molecular-probes-the-handbook.html [Accessed April 21, 2017].

50. Winkelman JD, Bilancia CG, Peifer M, Kovar DR (2014) Ena/VASP Enabled is a highly processive actin polymerase tailored to self-assemble parallel-bundled F-actin networks with Fascin. Proc Natl Acad Sci 111(11):4121–4126.

51. Jönsson P, Jonsson MP, Tegenfeldt JO, Höök F (2008) A Method Improving the Accuracy of Fluorescence Recovery after Photobleaching Analysis. Biophys J 95(11):5334–5348.

52. Shimozawa T, et al. (2013) Improving spinning disk confocal microscopy by preventing pinhole cross-talk for intravital imaging. Proc Natl Acad Sci 110(9):3399–3404.

53. Pubchem L-glutamic acid | C5H9NO4 - PubChem. Available at: https://pubchem.ncbi.nlm.nih.gov/compound/L-glutamic_acid [Accessed May 8, 2017].

54. Spruijt E, Westphal AH, Borst JW, Cohen Stuart MA, van der Gucht J (2010) Binodal Compositions of Polyelectrolyte Complexes. Macromolecules 43(15):6476–6484.

55. Hanke F, Serr A, Kreuzer HJ, Netz RR (2010) Stretching single polypeptides: The effect of rotational constraints in the backbone. EPL Europhys Lett 92(5):53001.

